# Early Archean origin of Photosystem II

**DOI:** 10.1101/109447

**Authors:** Tanai Cardona, Patricia Sánchez-Baracaldo, A. William Rutherford, Anthony W. D. Larkum

## Abstract

Photosystem II is a photochemical reaction center that catalyzes the light-driven oxidation of water to molecular oxygen. Water oxidation is the distinctive photochemical reaction that permitted the evolution of oxygenic photosynthesis and the eventual rise of Eukaryotes. At what point during the history of life an ancestral photosystem evolved the capacity to oxidize water still remains unknown. Here we study the evolution of the core reaction center proteins of Photosystem II using sequence and structural comparisons in combination with Bayesian relaxed molecular clocks. Our results indicate that a homodimeric photosystem with sufficient oxidizing power to split water had already appeared in the early Archean about a billion years before the most recent common ancestor of all described Cyanobacteria capable of oxygenic photosynthesis, and well before the diversification of some of the known groups of anoxygenic photosynthetic bacteria. Based on a structural and functional rationale we hypothesize that this early Archean photosystem was capable of water oxidation and had already evolved some level of protection against the formation of reactive oxygen species, which would place primordial forms of oxygenic photosynthesis at a very early stage in the evolutionary history of life.

## Introduction

The transition from anoxygenic to oxygenic photosynthesis initiated when an ancestral photochemical reaction center evolved the capacity to oxidize water to oxygen (Rutherford, 1989). Today, water oxidation is catalyzed in the Mn_4_CaO_5_ oxygen-evolving cluster of Photosystem II (PSII) of Cyanobacteria and photosynthetic eukaryotes. How and when Type II reaction centers diversified, and how and when one of these reaction centers evolved the capacity to oxidize water are questions that still remain to be answered. While there is agreement that by 3.5 Ga (billion years before the present) a form of anoxygenic photoautotrophy had already evolved (Tice & Lowe, 2004; Nisbet & Fowler, 2014; Butterfield, 2015), the sedimentological and isotopic evidence for the evolution of oxygenic photosynthesis ranges from 3.7 (Rosing & Frei, 2004; Frei et al., 2016) to the Great Oxidation Event (GOE) at ~2.4 Ga (J. E. Johnson et al., 2013). Similarly, molecular clock studies have generated a wider range of age estimates for the origin of Cyanobacteria between 3.5 (Falcon et al., 2010) to 2.0 Ga (Shih, Hemp, et al., 2017). There is thus great uncertainty and no consensus. For this reason, determining when Photosystem II evolved the capacity to oxidize water should greatly advance our understanding of the origin of oxygenic photosynthesis.

The evolution of Type II reaction center proteins has been described and discussed in some detail before (Beanland, 1990; Nitschke & Rutherford, 1991; Blankenship, 1992; Rutherford & Nitschke, 1996; Sadekar et al., 2006; Cardona, 2015, 2016) and it is presented and schematized in Fig. 1. Type II reaction centers can be divided in two major families: the *oxygenic* and the *anoxygenic* Type II reaction centers. The oxygenic Type II reaction center is also known as Photosystem II, and its electron transfer core is made of two homologous reaction center proteins, D1 and D2, exclusively found in Cyanobacteria and photosynthetic eukaryotes. The Mn_4_CaO_5_ cluster is bound by D1 and the core antenna protein CP43 (Ferreira et al., 2004). On the other hand, anoxygenic Type II reaction centers are found in phototrophic members of the phyla Proteobacteria, Chloroflexi, and Gemmatimonadetes, with the latter obtaining the reaction center via horizontal gene transfer from a gammaproteobacterium (Zeng et al., 2014). The core subunits of the anoxygenic Type II reaction centers are known as L and M and lack an oxygen-evolving cluster.

**Figure 1.**
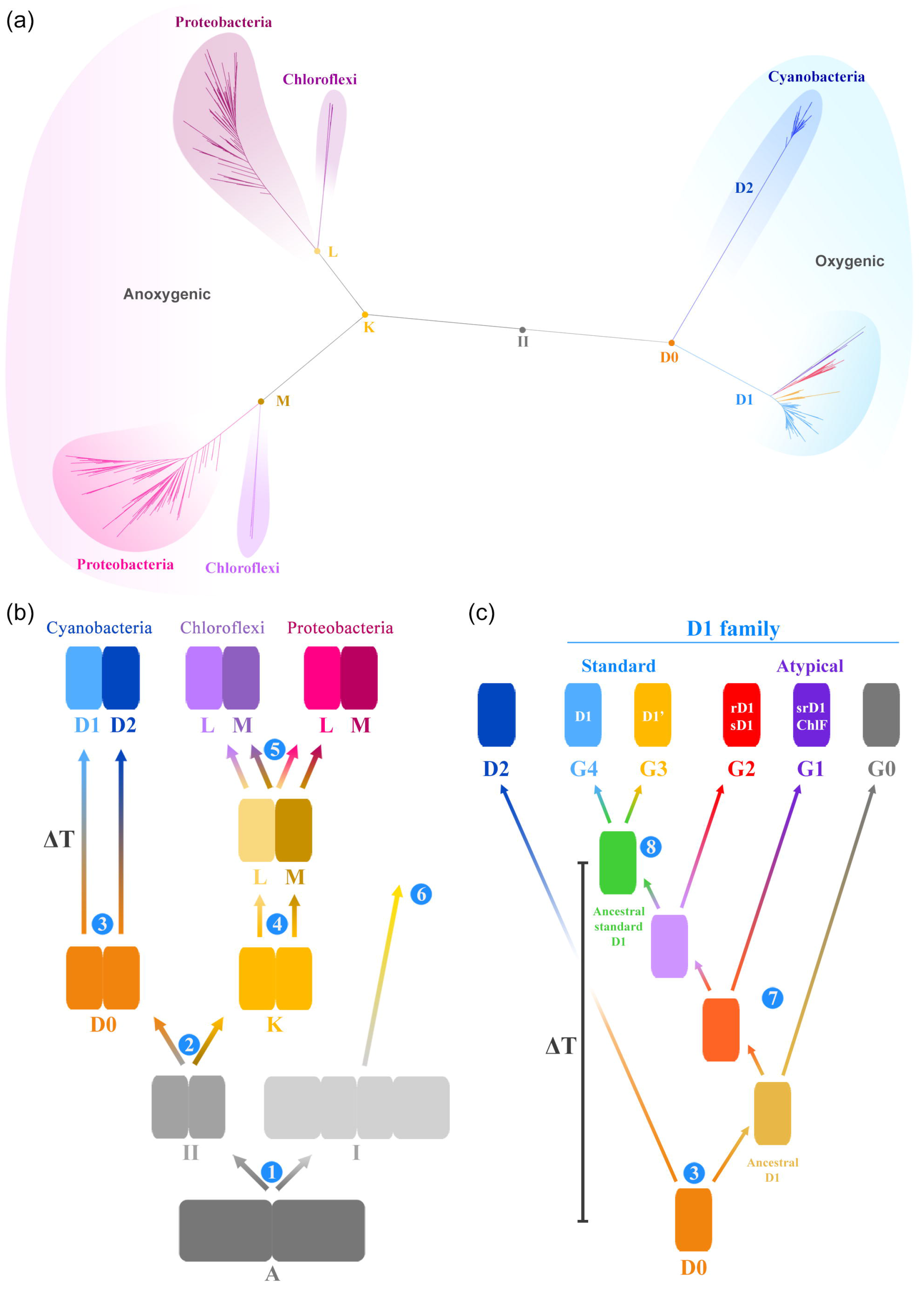
Evolution of Type II reaction center proteins. (a) A Maximum Likelihood phylogeny of Type II reaction center proteins. (b) A schematic representation of the phylogeny shown in (a). All reaction centers have common ancestry and descended from a homodimeric reaction center, marked **A**. From **A**, two new reaction centers emerged, one ancestral to all Type II reaction centers (**II**) and a second ancestral to all Type I reaction centers. This is the earliest diversification event of reaction center proteins that can be inferred from sequence and structural data and it is marked **1**. The ancestral Type II reaction center protein (**II**) gave rise to two new proteins, one ancestral to D1 and D2, named here **D0** and a second ancestral to L and M named **K**. The ancestral L and M subunits further diversify into Chloroflexi-type and Proteobacteria-type L and M subunits (5). Step **6** indicates that Type I reaction center proteins also diversified in parallel to Type II reaction center proteins. (c) Evolution of cyanobacterial D1 and D2, modified from Cardona et al. (2015). G0, G1, and G2 represent atypical D1 forms, and G3 and G4 standard D1 forms. **ΔT** marks the span of time between **D0** and the appearance of the ancestral standard form of D1, which characterizes Photosystem II and predates the most recent common ancestor of all known Cyanobacteria capable of oxygenic photosynthesis

There is no doubt that D1, D2, L, and M, share a common origin: Beanland (1990) was the first to record this but has been followed by many others (Nitschke & Rutherford, 1991; Sadekar et al., 2006; Cardona, 2015). That is to say that D1, D2, L and M, all descended from a single protein (denoted **II** in Fig. 1). The earliest event in the evolution of Type II reaction centers can be described as the divergence of this ancestral protein into two new forms, one ancestral to D1 and D2, the *oxygenic* branch; and a second one ancestral to L and M, the *anoxygenic* branch (Fig. 1). Hence, D1 and D2 originated from a gene duplication event and together make a monophyletic clade of Type II reaction center proteins, distinct from that which gave rise to L and M (Cardona, 2015, 2016b). The ancestral protein to D1 and D2 will be referred to as **D0** and the ancestral protein to L and M will be referred to as **K**.

As a result of the monophyletic relationship of D1 and D2 and the conserved structural and functional characteristics between these two proteins, it is possible to reconstruct traits of the ancestral photosystem. Some of the conserved traits, present in both D1 and D2, but absent in L and M, suggest that the ancestral homodimeric photosystem, made of a **D0** dimer, was already unlike any of the known anoxygenic Type II reaction centers and had acquired characteristics associated with the highly oxidizing potential required for water oxidation (Rutherford & Nitschke, 1996; Rutherford & Faller, 2003; Cardona et al., 2012; Cardona, 2016). One of these conserved traits is a redox tyrosine-histidine pair strictly conserved in both D1 and D2, Y_Z_-H190 and Y_D_-H189, respectively. The presence of these tyrosine-histidine pairs indicates that the midpoint potential (E_*m*_) of the photochemical chlorophylls at the heart of the reaction center was oxidizing enough to generate the neutral tyrosyl radical on either side of the homodimeric reaction center (Rutherford & Nitschke, 1996; Rutherford & Faller, 2003). That is an E_*m*_ of at least 1 V (DeFelippis et al., 1989; DeFelippis et al., 1991), sufficient to drive the oxidation of water to oxygen, which has an E_*m*_ of 0.82 V (Dau & Zaharieva, 2009; Tachibana et al., 2012). Based on this and other arguments, Rutherford and Nitschke (1996) suggested that before the gene duplication that led to D1 and D2 this ancestral photosystem was well on its way towards the evolution of water oxidation, and may have been able to oxidize water, even if only inefficiently.

Several types of D1 can be distinguished phylogenetically (Cardona et al., 2015) and their evolution is schematized in Fig. 1C. The early evolving forms, referred to as atypical D1 forms (G0, G1, G2 in Fig. 1), are characterized by the absence of some, but not all of the ligands to the Mn_4_CaO_5_ cluster and have been recently found to be involved in the synthesis of chlorophyll *f*, which supports oxygenic photosynthesis using low energy far-red light (M. Y. Ho et al., 2016); or the inactivation of PSII when anaerobic process are being carried out such as nitrogen fixation (Murray, 2012; Wegener et al., 2015). The late evolving forms, referred to as the standard D1 forms, are characterized by a complete set of ligands to the Mn_4_CaO_5_ cluster and are the main D1 used for water oxidation. Among the standard forms there are also several types, which have been roughly subdivided in two groups: the microaerobic forms of D1 (G3) and the dominant form of D1 (G4). The microaerobic forms are suspected to be expressed only under low-oxygen conditions. The dominant form, G4, is the main D1 used for water oxidation by all Cyanobacteria and photosynthetic eukaryotes. Most Cyanobacteria carry in their genomes an array of different D1 types, yet every strain has at least one dominant form of D1 (G4). Therefore, all Cyanobacteria descended from a common ancestor that already had evolved efficient oxygenic photosynthesis, had a dominant form of D1, and was able to assemble a standard Photosystem II virtually indistinguishable from that of later evolving strains. Furthermore, because the atypical D1 forms support or regulate oxygenic photosynthesis under specific environmental conditions it can be argued that when these branched out water oxidation to oxygen had already evolved.

Based on the phylogeny of reaction center proteins several stages in the evolution of oxygenic photosynthesis can be envisaged: the earliest of these stages is the divergence of Type I and Type II reaction center proteins (**1**, Fig. 1B); this is then followed by the divergence of the *anoxygenic* family (L/M) and the *oxygenic* family (D1/D2) of Type II reaction center proteins (**2**), then by the duplication event that led to the divergence of D1 and D2 (**3**), and the subsequent (**7**) gene duplication events and specializations that created the known diversity of D1 forms, which ultimately resulted in the emergence of the standard form of D1. Because a photosystem made of a **D0** had already acquired some of the fundamental features required to oxidize water such as a highly oxidizing chlorophyll cofactors and the capacity to generate the neutral tyrosyl radical at each side of the reaction center: then it can be suggested that some of the earliest stages specific to the evolution of Photosystem II and oxygenic photosynthesis had occurred between stages **2** and **3** as depicted in Fig. 1B. Therefore, the span of time between **D0** and the ancestral standard form of D1 (marked **8** in Fig. 1C) represents the duration of the evolutionary trajectory of Photosystem II from a simpler homodimeric highly-oxidizing reaction center to the more complex enzyme inherited by all organisms capable of oxygenic photosynthesis. We denote this span of time by **ΔT**. If **ΔT** is small, such as a few million years or less for example, then the evolution of oxygenic photosynthesis may be better described as a sudden and fast process only getting started shortly before the GOE as suggested by some recent analyses (Ward et al., 2016; Shih, Hemp, et al., 2017). On the other hand, if **ΔT** is large: several hundred million years or more for example, then the earliest stages in the evolution of oxygenic photosynthesis could significantly predate the GOE as suggested by some geochemical (Mukhopadhyay et al., 2014; Planavsky et al., 2014; Satkoski et al., 2015) and phylogenetic data (Blank & Sanchez-Baracaldo, 2010; Schirrmeister et al., 2013).

Here we report an in-depth evolutionary analysis of Type II reaction center proteins including Bayesian relaxed molecular clocks under various scenarios for the origin of photosynthesis. The data presented here indicate that a photosystem with the structural and functional requirements for the evolution of water oxidation could have arisen in the early Archean and long before the most recent common ancestor of Cyanobacteria.

## Results

### Change in sequence identity as a function of time

A first approximation to the evolution of Type II reaction centers as a function of time can be derived from the level of sequence identity between D1 and D2 of different species with known or approximated divergence times as shown in Fig. 2. For example, the D1 protein of the dicotyledon *Arabidopsis thaliana* shares 99.7% amino acid sequence identity with that of the dicotyledon *Populus trichocarpa,* and these are estimated to have diverged between 127.2 and 82.8 Ma (Clarke et al., 2011), see Fig. 2 and supplementary Table S1. On the other hand, *A. thaliana*’s D1 shares 87.7% sequence identity with that of a unicellular red alga *Cyanidioschyzon merolae*. Complex multicellular red algae are known to have diverged at least 1.0 Ga ago (Butterfield, 2000; Gibson et al., 2017) and recently described fossils could push this date back to 1.6 Ga (Bengtson et al., 2017; Sallstedt et al., 2018). At the other end of this evolutionary line, the three dominant forms of D1 from *Gloeobacter violaceous* share on average 79.2% sequence identity with that of *C. merolae* or 78.5% with that of *A. thaliana*. If the percentage of sequence identity between pairs of species is plotted as a function of their divergence time, a linear decrease of identity is observed among reaction center proteins at a rate of less than 1% per 100 million years (supplementary Table S2). The trend in Fig. 2 indicates that the rate of evolution of D1 and D2 since the GOE and since the emergence of photosynthetic eukaryotes has remained relatively slow until the present time, if considered over a large geological time scale, with less than 20% change in sequence identity in the past 2.0 Ga.

**Figure 2.**
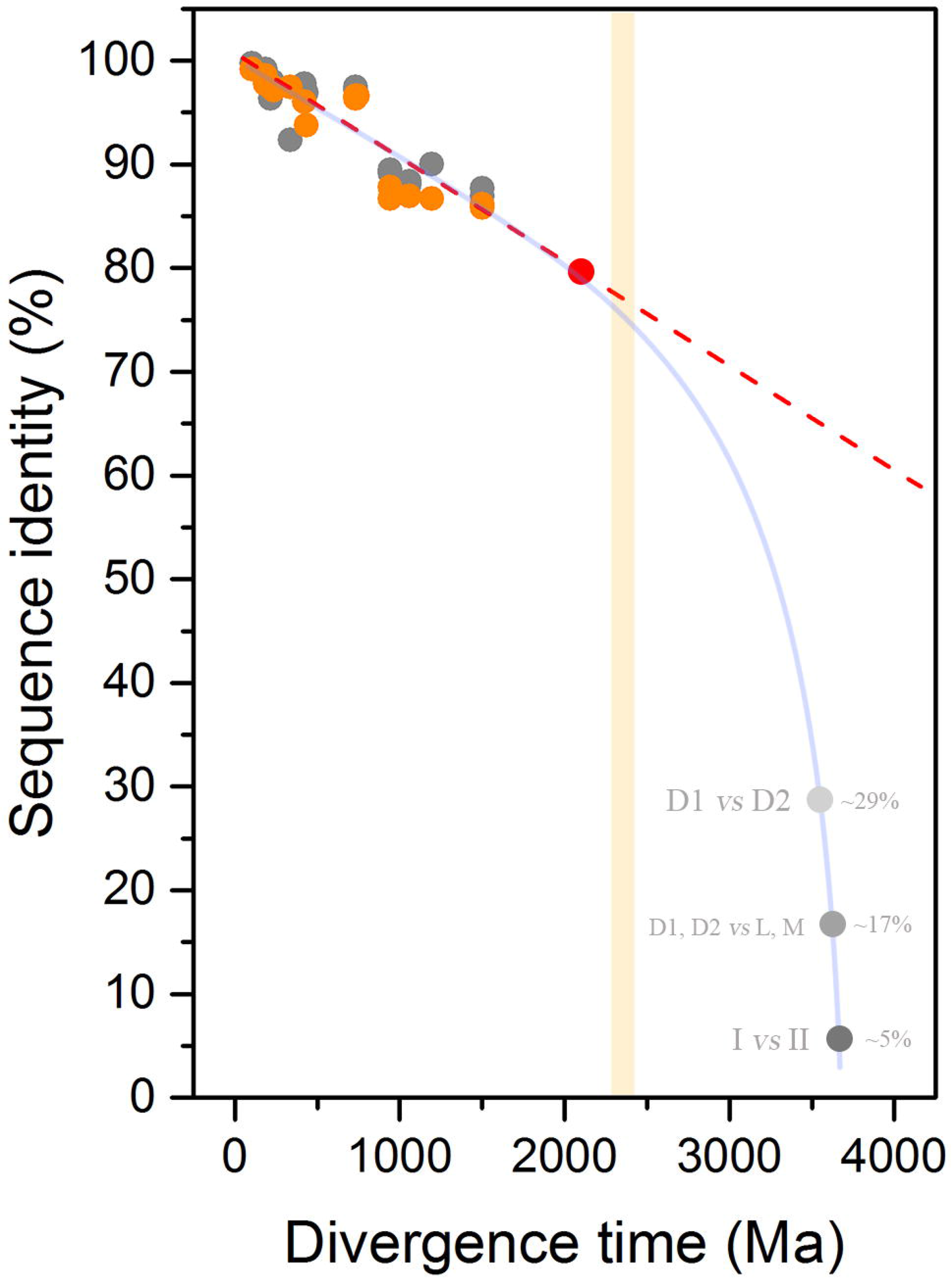
Decrease of sequence identity of D1 and D2 proteins as a function of divergence time. D1 subunits are shown in gray and D2 in orange. The divergence time between pairs of species is plotted against the level of sequence identity as tabulated in supplementary Table S1. The red circle, placed at 79.2% corresponds to the average sequence identity of the three distinct Group 4 D1 sequences of *Gloeobacter violaceous* in comparison with that of *Cyanidioschyzon merolae*. The light orange bar marks the GOE. The dashed line is fitted from a linear function and shows that over a period of at least 2.0 Ga no dramatic changes in the rates of evolution of D1 and D2 are observed. The red dashed lines shows an extrapolation of current rates of evolution throughout Earth’s history. This line highlights that the rate is too slow for the divergence of D1 and D2 to have started right before the GOE. The gray dots around 3.5-3.8 Ga marks an especulative timing for the earliest events in the history of photosynthesis: the divergence of D1 and D2 (~29% sequence identity), the divergence of anoxygenic (L/M) and oxygenic (D1/D2) reaction center proteins (~17%), and the divergence of Type I and Type II reaction center proteins (≤5%). The curved blue line highlights that any scenario for the diversification of reaction centers after the origin of life requires faster rates of evolution at the earliest stages in the evolution of photosynthesis

Now, if the most recent common ancestor (MRCA) of Cyanobacteria, defined as the MRCA of the genus *Gloeobacter* and all other extant photosynthetic strains, existed hundreds of millions of years before the GOE, this would presuppose an even slower rate of evolution of the core subunits of PSII. On the other hand, if the rate of evolution of D1 and D2 are taken at face value, following the roughly uniform rate observed in photosynthetic eukaryotes, this would locate the divergence of *Gloeobacter* after the GOE (Fig. 2, red spot): in consequence, the older the MRCA of Cyanobacteria, the slower the rate of evolution of the dominant form of D1 and D2. Therefore, large uncertainties in the fossil record of photosynthetic eukaryotes would result in only small changes to this trend. For example, if the divergence of red algae occurred as late as 1.0 Ga or as early as 2.0 Ga, this will only cause a small shift in the overall rate. Or for example, if the MRCA of angiosperms is actually 100 million years older than currently understood, this would result in almost a negligible change in the rate of evolution of the dominant form of D1 and D2 compared over the large time scale of the planet.

Let us reiterate that all the evidence suggests that all reaction center proteins originated from a single ancestral protein that diversified as the multiple groups of photosynthetic bacteria arose. As a result of this common ancestry, any standard D1 shares on average about 29% sequence identity with any D2 across their entire sequence. Any standard D1 or D2 share on average 17% sequence identity with any L or M. The level of sequence identity falls well below 5% if any Type II reaction center protein is compared with any Type I reaction center protein (Cardona, 2015). As a result of this, the rate of evolution of D1 and D2 since the GOE, as estimated from the decrease of sequence identity (<1% per 0.1 Ga), is too slow to account for the evolution of photochemical reaction centers within a reasonable amount of time (Fig. 2, dashed line). In other words, the rate of evolution of reaction center proteins since the origin of life could not have been constant, and any scenario for the origin of photochemical reaction centers at any point in the Archean requires initially faster rates of evolution than any rate observed since the Proterozoic (Fig. 2, light blue line).

Taking into consideration that D1 and D2 share only about 29% sequence identity, this would suggest, as illustrated in Fig. 2, that the event that led to the divergence of D1 and D2 is more likely to have occurred closer to the origin of the primordial reaction center proteins during the emergence of the first forms of photoautotrophy in the early Archean, than closer to or after the GOE.

### Bayesian relaxed molecular clock analysis

The simple approach used above indicates that the divergence of D1 and D2 is likely placed well before the GOE, to confirm this observation we applied a molecular clock to the phylogeny of Type II reaction center proteins. Figure 3 shows a Bayesian relaxed log-normal autocorrelated molecular clock built using the CAT + GTR + Γ model allowing for flexible boundaries on the calibration points (Lartillot et al., 2009). As an informed starting point we first specified the age of the root (root prior) at 3.5 Ga with a standard deviation of 0.05 Ga. That is to say, that the most ancestral form of a Type II reaction center protein is assumed to have already evolved by 3.5 Ga. Under these conditions, the last common ancestral protein to the standard form of D1 prior to the divergence of the G3 and G4 types (Fig. 3, green dot) is timed at 2.19 ± 0.22 Ga. On the other hand, **D0** (Fig. 3, orange dot) is timed at 3.22 ± 0.19 Ga. It follows then that the difference in time between **D0** and the first standard form of D1, **ΔT**, is 1.02 Ga, with the level of uncertainty on the estimated ages resulting in a range for **ΔT** between 1.44 to 0.60 Ga (see Table 1 and Fig. 4 A and B). This large **ΔT** is in agreement with the inferences made from the comparisons of sequence identity plotted in Fig. 2.

**Figure 3.**
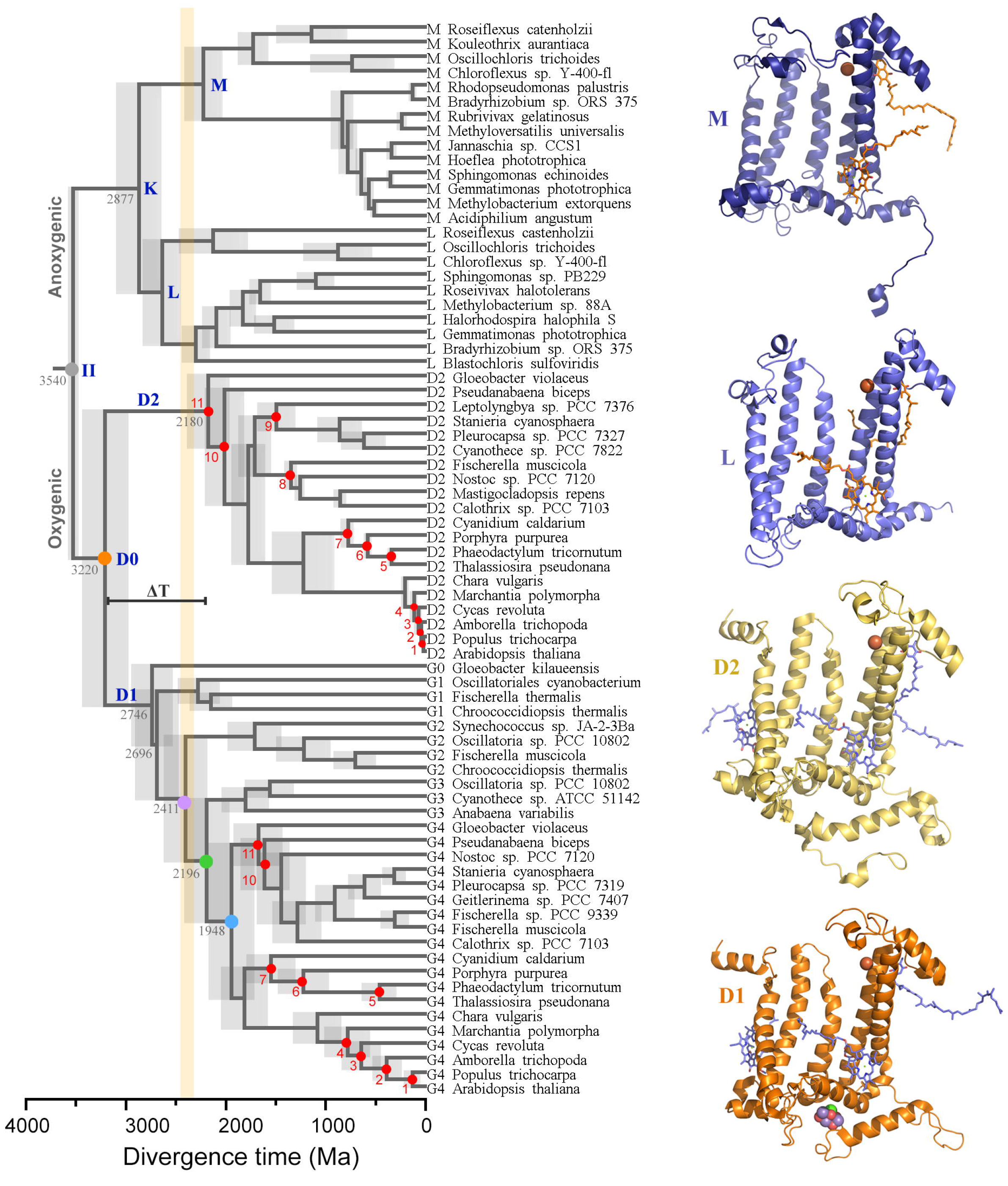
Relaxed molecular clock of Type II reaction center proteins. A log-normal autocorrelated relaxed clock is shown implementing a CAT + GTR + Γ non-parametric model with flexible boundaries on the calibration points. Red dots are calibration points as described in Materials and Methods. The gray dot denoted **II**, represents the ancestral Type II reaction center protein, as schematized in Fig. 1. The orange dot (**D0**) marks the initial divergence of D1 and D2. The violet dot marks the divergence point between G2 atypical D1 sequences and standard D1. The green dot marks the divergence point between the microaerobic D1 forms (G3) and the dominant form of D1 (G4). This point represents the last common ancestral protein to all standard D1 forms predating crown group Cyanobacteria. The blue dot represents the origin of the dominant form of D1 inherited by all extant Cyanobacteria and photosynthetic eukaryotes. The gray bars represent the standard error of the estimated divergence times at the nodes. The orange bar shows the GOE

**Table 1.** Effect on **ΔT** assuming different ages for the most ancestral Type II reaction center protein

| Root prior<br>(Ga) | II | D0 | Ancestral<br>standard D1 | $\Delta T$ (range) |
| --- | --- | --- | --- | --- |
| CAT + GTR + $\Gamma$ | | | | |
| **3.2** | $3.25 \pm 0.05$ | $2.80 \pm 0.16$ | $1.99 \pm 0.19$ | 0.80 (1.17 - 0.44) |
| **3.5** | $3.54 \pm 0.05$ | $3.22 \pm 0.19$ | $2.19 \pm 0.24$ | 1.02 (1.44 - 0.60) |
| **3.8** | $3.83 \pm 0.05$ | $3.44 \pm 0.21$ | $2.27 \pm 0.24$ | 1.17 (1.62 - 0.71) |
| **4.1** | $4.12 \pm 0.05$ | $3.71 \pm 0.23$ | $2.38 \pm 0.25$ | 1.32 (1.81 - 0.84) |
| CAT + GTR + $\Gamma$ and removing calibration point 11 | | | | |
| **3.5** | $3.52 \pm 0.05$ | $3.00 \pm 0.29$ | $1.78 \pm 0.25$ | 1.22 (1.77 - 0.66) |
| **3.8** | $3.81 \pm 0.05$ | $3.15 \pm 0.29$ | $1.77 \pm 0.25$ | 1.37 (1.93 - 0.81) |
| LG + $\Gamma$ | | | | |
| **3.2** | $3.27 \pm 0.05$ | $3.19 \pm 0.08$ | $2.51 \pm 0.13$ | 0.68 (0.89 - 0.46) |
| **3.5** | $3.53 \pm 0.05$ | $3.40 \pm 0.09$ | $2.64 \pm 0.15$ | 0.77 (1.00 - 0.53) |
| **3.8** | $3.81 \pm 0.05$ | $3.64 \pm 0.12$ | $2.77 \pm 0.17$ | 0.88 (1.16 - 0.58) |
| **4.1** | $4.10 \pm 0.05$ | $3.91 \pm 0.14$ | $2.90 \pm 0.19$ | 1.01 (1.34 - 0.68) |
| LG + $\Gamma$ and removing calibration point 11 | | | | |
| **3.5** | $3.49 \pm 0.05$ | $3.18 \pm 0.19$ | $2.30 \pm 0.20$ | 0.87 (1.26 - 0.48) |
| **3.8** | $3.79 \pm 0.05$ | $3.52 \pm 0.19$ | $2.55 \pm 0.22$ | 1.38 (0.97 - 0.55) |

**Figure 4.**
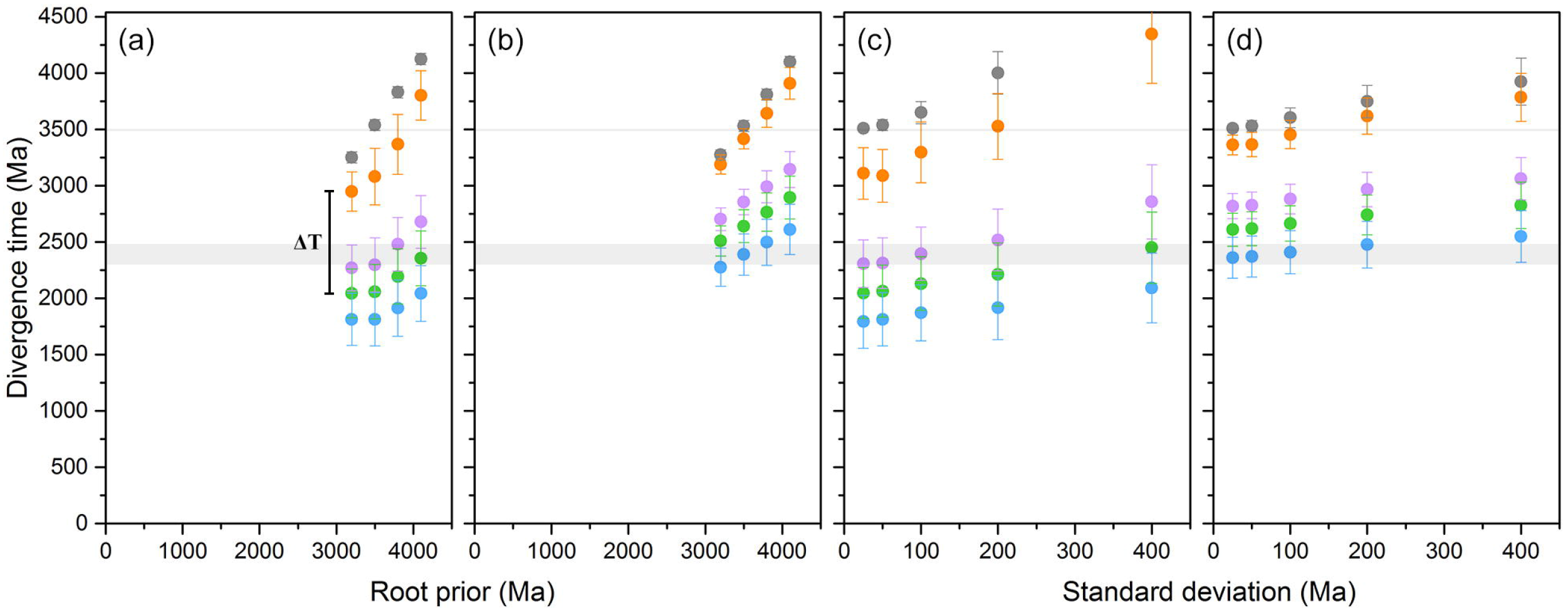
Effect of model selection on estimated divergence times. (*a*) Divergence times of key nodes in the evolution of Type II reaction centers as a function of the root prior. The root prior was varied from 3.2 to 4.1 ± 0.05 Ga under a CAT + GTR + Γ model. The colored dots match selected nodes of interest in Fig. 3. The thick gray bar marks the GOE and the narrow bar marks the minimum accepted age for the origin of photosynthesis. (*b*) Identical to (*a*) but using a LG + Γ model of amino acid substitutions. (*c*) Divergence times of key nodes assuming a root prior of 3.5 Ga as a function of the standard deviation on the root. The standard deviation was varied from 0.025 to 0.4 Ga under a CAT + GTR + Γ model. (*d*) Identical to (*c*) but using a LG + Γ model. In every case **D0** is the oldest node after the root and the magnitude of **ΔT** is always in the range of a billion years

To test the effect of different root priors on our results we varied the age of the root and the standard deviation over a broad range. Table 1 lists estimates of divergence times of key ancestral Type II reaction center proteins and the respective **ΔT** value using different root priors. For example, under the assumption that Type II reaction centers had already evolved by 3.8 Ga ago (Rosing, 1999; Czaja et al., 2013; Nisbet & Fowler, 2014), **ΔT** is found to be centered at 1.17 Ga, with a range between 1.62 and 0.71 Ga. Similarly, if it is assumed to be a late event occurring at 3.2 Ga, though unlikely, **ΔT** is still 0.80 Ga with a range between 1.17 and 0.44 Ga. Furthermore, increasing the standard deviation on the root prior pushes the timing of the earliest events in the evolution of Type II reaction centers to even older ages rather than younger ages, see Table 2 and Fig. 4 C and D. For example, a root prior of 3.5 Ga with a standard deviation of 0.1 Ga pushes the estimated time for the root to 3.65 Ga; making **D0** 3.30 ± 0.27 and generating a **ΔT** of over a billion years.

**Table 2.** Effect of varying the standard deviation (S. D.) on the root prior at 3.5 Ga

| **S. D.**<br>**(Ga)** | **II** | **D0** | **Ancestral**<br>**standard D1** | **$\Delta T$ (range)** |
| --- | --- | --- | --- | --- |
| **CAT + GTR + $\Gamma$** |  |  |  |  |
| **0.025** | $3.50 \pm 0.03$ | $2.99 \pm 0.22$ | $2.07 \pm 0.22$ | 0.91 (1.31 - 0.46) |
| **0.05** | $3.54 \pm 0.05$ | $3.22 \pm 0.19$ | $2.19 \pm 0.24$ | 1.02 (1.44 - 0.60) |
| **0.10** | $3.65 \pm 0.10$ | $3.29 \pm 0.26$ | $2.22 \pm 0.23$ | 1.07 (1.56 - 0.57) |
| **0.20** | $4.00 \pm 0.19$ | $3.61 \pm 0.32$ | $2.32 \pm 0.25$ | 1.32 (1.87 - 0.70) |
| **0.40** | $4.82 \pm 0.38$ | $4.55 \pm 0.44$ | $2.50 \pm 0.28$ | 1.74 (2.48 - 1.01) |
| **LG + $\Gamma$** |  |  |  |  |
| **0.025** | $3.51 \pm 0.02$ | $3.36 \pm 0.09$ | $2.61 \pm 0.15$ | 0.75 (0.98 - 0.51) |
| **0.05** | $3.53 \pm 0.05$ | $3.37 \pm 0.11$ | $2.62 \pm 0.15$ | 0.75 (1.00 - 0.48) |
| **0.10** | $3.60 \pm 0.09$ | $3.45 \pm 0.13$ | $2.67 \pm 0.16$ | 0.79 (1.07 - 0.50) |
| **0.20** | $3.75 \pm 0.14$ | $3.62 \pm 0.16$ | $2.74 \pm 0.18$ | 0.88 (1.21 - 0.53) |
| **0.40** | $3.92 \pm 0.21$ | $3.79 \pm 0.21$ | $2.83 \pm 0.21$ | 0.96 (1.37 - 0.53) |

In Fig. 3 it can also be seen that the divergence of the L and M subunits occurs *after* the divergence of D1 and D2. The estimated time for the divergence of L and M is 2.87 ± 0.16 Ga, while the time for the divergence of D1 and D2, as we saw above, is 3.22 ± 0.19 Ga. The late divergence of L and M suggests that the divergence of D1 and D2 predates significantly the diversification event that led to the evolution of Chloroflexi-type and Proteobacteria-type L and M subunits. These results confirm that Cyanobacteria did not obtain Type II reaction centers via horizontal gene transfer (HGT) of L and M from an early evolving representative of the phylum Chloroflexi or Proteobacteria (Cardona, 2015).

The Bayesian clock using flexible boundaries on the calibration points consistently produced ages for the divergence of the D2 subunit of *G. violaceous* and the dominant form of D1 (G4) after the GOE, similar to the ages reported by Shih, Hemp, et al. (2017). Yet, previous molecular clocks have suggested that the MRCA of Cyanobacteria might predate the GOE (Schirrmeister et al., 2015), so we also performed a similar analysis that allowed us to explore this scenario. This was achieved using an empirical amino acid substitution model (LG + Γ) instead of the non-parametric approach described above. We found this to be the only way to locate the D2 of *G. violaceous* and the dominant form of D1 (G4) before the GOE. The effect of less flexible boundaries on the estimated divergence times is shown in Table 1 and Table 2, and Fig. 4 B and D. For example, assuming a root at 3.5 ± 0.05 Ga, the estimated divergence time for the standard form of D1 becomes 2.64 ± 0.15 Ga and pushes **D0** back to 3.40 ± 0.09 Ga, making **ΔT** 0.77 Ga with a range between 1.00 and 0.53 Ga. On the other hand, if we allowed flexibility on the root prior by increasing the standard deviation to 0.4 Ga (Table 2), the estimated divergence time for the standard form of D1 becomes 2.83 ± 0.21, but the estimated age of the root is pushed back to 3.92 ± 0.21 Ga with **D0** at 3.79 ± 0.21, making **ΔT** about a billion years. Overall, placing the MRCA of Cyanobacteria before the GOE pushes the gene duplication event that led to the divergence of D1 and D2 even closer to the origin of Type II reaction centers and to the origin of photosynthesis.

### Rates of evolution

The inferences derived from Fig. 2 revealed that the rates of evolution had to be faster in the initial stages during the Archean compared with the Proterozoic, even when **ΔT** is as large as one billion years. To gain a better understanding of the changes of the rate of evolution of Type II reaction center proteins we plotted the rates as a function of divergence time. In Fig. 5A the rate of evolution (**ν**) of each node in the tree, expressed as amino acid substitutions per site per unit of time, is plotted against the estimated divergence time for each respective node. It can be seen that the rate at the earliest stage is much faster than the rates observed since the Proterozoic. Thus, faster rates are necessary to explain the origin and evolution of Type II reaction centers at any point in the Archean and regardless of when exactly photosynthesis originated, as seen in Fig. 5B. The decrease in the rate of evolution is consistent with the observations derived from Fig. 2 and can be roughly fitted with a first-order exponential decay curve (fitting parameters are presented in supplementary Table S3). Fig. 5A additionally shows that L and M have been evolving at a slightly faster rate than D1 and D2. From this slow-down of the rates it can be calculated that since each respective duplication event (stages **2** and **4** in Fig. 1) it took about 168 million years for D1 and D2 to fall to 50% sequence identity and about 115 million years for the same to occur for L and M

**Figure 5.**
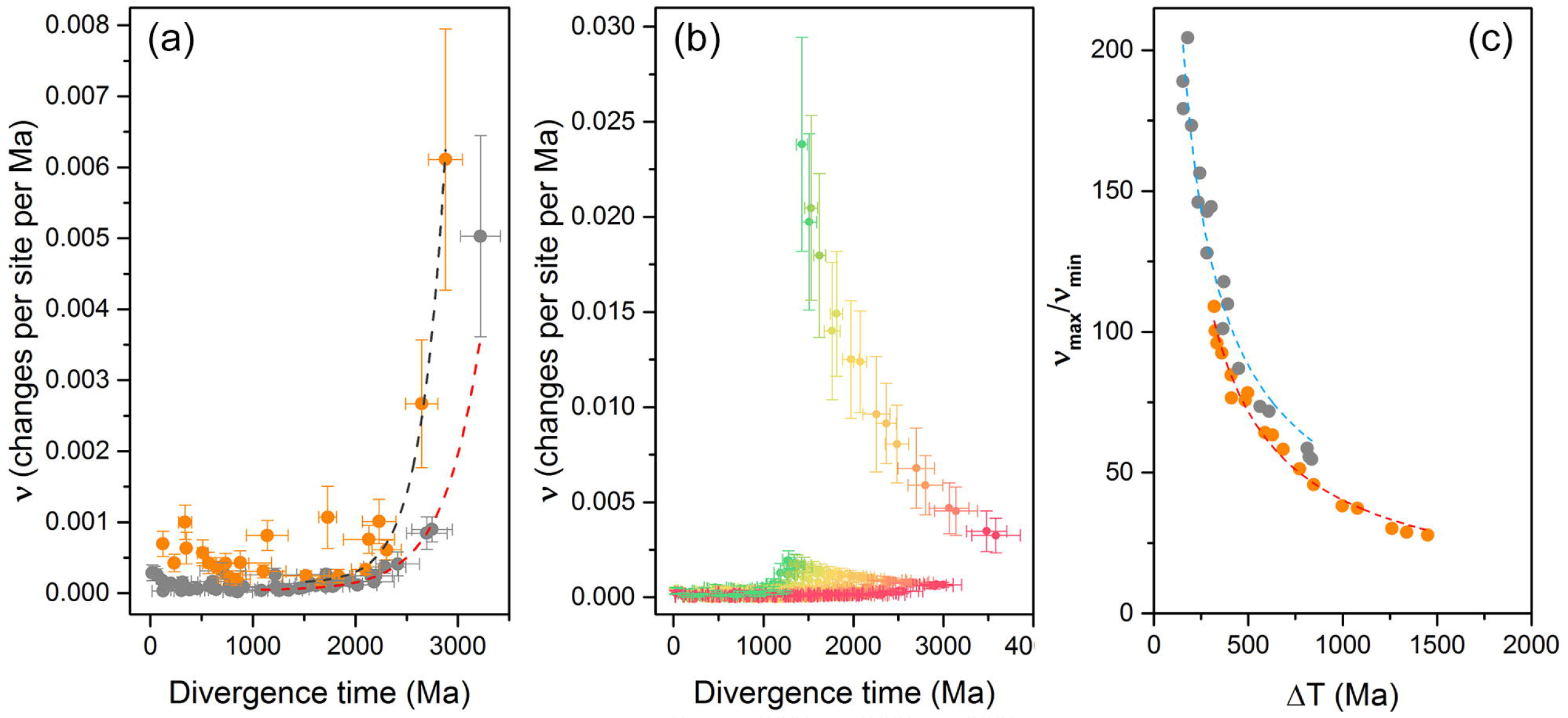
Rates of evolution as a function of time. (*a*) Change in the rate of evolution of oxygenic (gray) and anoxygenic (orange) Type II reaction center proteins. The rates correspond to the tree in Fig. 3, assuming an origin of photosynthesis at about 3.5 Ga. The dashed lines represent a fit of a single-component exponential decay and the rates are given as amino acid substitutions per site per million years. (*b*) Changes in the rate of evolution constraining the root to younger and younger ages. The red curve farthest to the right was calculated using a root prior of 4.2 ± 0.05 Ga, while the green curve farthest to the left was calculated using a root prior of 0.8 ± 0.05 Ga. Younger divergence times implies initial faster rates of evolution. (*c*) Change in the rate of evolution as a function of **ΔT**, the dashed lines represents a fit to a power-law function. The curve in orange was calculated using **ΔT** values subtracting the mean average of the divergence times of **D0** and the ancestral standard D1. The curve in gray was calculated using **ΔT** values subtracting the minimum age of **D0** and the maximum age for ancestral standard D1

The maximum rate of evolution in the D1 and D2 family of reaction center proteins is placed at the node that represents **D0**. We will refer to this rate as **ν**_**max**_. Fig. 5A shows that the rate of evolution flattens out to comparatively slow rates during the Proterozoic. These rates correspond to the rates of Group 4 D1 and D2. We will refer to the average rate of evolution during this zone of slow change as **ν**_**min**_ and it is calculated as the average rate from each node in Group 4 D1 and D2. In Fig. 5A, **ν**_**max**_ is 5.03 ± 1.42 amino acid substitutions per site per Ga, while **ν**_**min**_ is 0.12 ± 0.04 substitutions per site per Ga. Therefore, if Type II reaction centers had evolved by 3.5 Ga, to account for the divergence of D1 and D2 in one billion years, the initial rate of evolution had to be about 40 times larger than that observed since the MRCA of Cyanobacteria.

Table 3 lists the rates of evolution of a diverse number of proteins reported in independent studies in a broad range of organisms. It is found that the rate of evolution of the core subunits of PSII (**ν**_**min**_) is similar to the rate of other proteins that are very ancient and highly conserved such as subunits of the ATPase, the Cytochrome *b*_*6*_*f* complex, or the ribosome. Our estimated rates fall well within the expected range of other cyanobacterial proteins, thus validating our calibration choices and consistent with the expected level of sequence identity as derived from Fig. 2.

**Table 3.** Comparison of rates of protein evolution

|  | **Rate**<br>**(Amino acid substitutions per**<br>**site per Ga)** | **References** |
| --- | --- | --- |
| D0 ($v_{\max}$) | 5.03 | This work<sup>a</sup> |
| Group 4 D1 and D2 ($v_{\min}$) | 0.12 | This work<sup>a</sup> |
| K | 6.11 | This work<sup>a</sup> |
| L and M | 0.40 | This work<sup>a</sup> |
| PsaA, Photosystem I core subunit<br>(Cyanobacteria) | 0.09 | (Sanchez-Baracaldo,<br>2015) |
| AtpA, ATP Synthase CF1 Alpha<br>Chain (Cyanobacteria) | 0.08 | (Sanchez-Baracaldo<br>2015) |
| PetB, Cytochrome *b6* (Cyanobacteria) | 0.05 | (Sanchez-Baracaldo<br>2015) |
| S13 Ribosomal Protein<br>(Cyanobacteria) | 0.13 | (Sanchez-Baracaldo<br>2015) |
| L1 Ribosomal Protein (Cyanobacteria) | 0.11 | (Sanchez-Baracaldo<br>2015) |
| ADP-glucose pyrophosphorylase large<br>subunit (plants) | 1.2 | (Georgelis et al.,<br>2008) |
| PRTB Protein<sup>b</sup> (humans) | 0.13 | (Matsunami et al.,<br>2011) |
| Alcohol dehydrogenase (ascidians) | 0.27 | (Canestro et al., 2002) |
| Protein-L-isoaspartyl (D-aspartyl) *O*-<br>methyltransferase (bacteria to<br>humans<sup>c</sup>) | 0.39 | (Kagan et al., 1997) |
| Hepatitis C Virus | 3700 | (Bukh et al., 2002) |
| Influenza virus type A (H1) | 5800 | (Carrat & Flahault,<br>2007) |

<sup>a</sup>Estimated using a root prior of 3.5 Ga under a autocorrelated log-normal molecular clock as described in the text and Material and Methods.
<sup>b</sup>Proline-rich transcript overexpressed in the brain (PRTB). The human protein shares 99% sequence identity compared to that in mice. Rodents are estimated to have diverged about 74 Ma ago (Kay & Hoekstra, 2008).
<sup>c</sup>The authors pointed out that the rate of evolution of this methyltransferase has remain unchanged from bacteria to humans.

Our Bayesian analyses suggests that the evolution of PSII is better described by a long span of time since the appearance of a homodimeric photosystem (with sufficient power to oxidize water) until the emergence of standard PSII (inherited by all known Cyanobacteria capable of photosynthesis). Notwithstanding, a fast rate of evolution at the earliest stage implies that **ν**_**max**_ would increase if **ΔT** is considered to be smaller, as would be the case for an evolutionary scenario in which PSII evolves rapidly before the GOE after an event of horizontal gene transfer of an anoxygenic Type II reaction center like those found in phototrophic Proteobacteria and Chloroflexi. We illustrate this concept in Fig. 5B and 5C. These figures depict the change of the rate of evolution as a function of **ΔT**. This manipulation of the molecular clock can only by accomplished computationally by changing the root prior to younger and younger ages. The increase of **ν**_**max**_ with decreasing **ΔT** can be fitted using a power law-function (supplementary Table S4). For example, in the hypothetical case that **ΔT** is 1 million years; that is, if the divergence of D1 and D2 occurred in only 1 million years, then **ν**_**max**_ would have been about 12500 amino acid changes per site per Ga, which is greater than the rate of evolution of some extant viruses, see Table 3. If **ΔT** is hypothesized to be 10 or 100 million years instead, this would require a **ν**_**max**_ of about 165 or 33 amino acid substitutions per site per Ga, respectively. A rate of about 33 substitutions per site per Ga means that 30 million years would be enough time to erase all sequence identity between D1 and D2, which is an implausible rate if photosynthesis originated in the early Archean (Tice & Lowe, 2004; Schopf et al., 2018) and for proteins that are, and may have always been, under strong purifying selection (Shi et al., 2005). Whether a highly conserved (slow evolving) core subunit of any protein complex in all bioenergetics can evolve, or has ever evolved, as such high rates is to the best of our knowledge unknown.

### Sensitivity analysis

To test the reliability of the method we explored a range of contrasting models. We compared the effect of the model of relative exchange rates on the estimated divergence times: Supplementary Fig. S1 provides a comparison of estimated divergence times calculated with the CAT + GTR model (Z. Yang & Rannala, 2006) against divergence times calculated using the CAT model with a uniform (Poisson) model of equilibrium frequencies. The GTR model does not have a strong effect on the calculated divergence times as the slope of the graph does not deviate from unity when paired with the uniform model (see supplementary Table S5 for linear regressions). Thus under a root prior of 3.5 ± 0.05 Ga a CAT + Poisson model also generated a **ΔT** centered at 1.02 Ga, see Table 4.

**Table 4.** Change in **ΔT** under different evolutionary models

| **Model** | **Root prior (Ga)** | **Calibration (Ga)** | **$\Delta T$ (Ga)** |
| --- | --- | --- | --- |
| CAT + GTR (autoc.<sup>a</sup>) | 3.5 | 2.45 | 1.02 (1.44 - 0.52) |
| CAT + GTR (autoc.) | 3.5 | 2.70 | 0.97 (1.29 - 0.64) |
| CAT + GTR (autoc.) | 3.8 | 2.45 | 1.19 (1.64 - 0.70) |
| CAT + GTR (autoc.) | 3.8 | 2.70 | 1.13 (1.48 - 0.76) |
| CAT + Pois. (autoc.) | 3.5 | 2.45 | 1.02 (1.51 - 0.52) |
| CAT + Pois. (autoc.) | 3.5 | 2.70 | 1.00 (1.34 - 0.66) |
| CAT + Pois. (autoc.) | 3.8 | 2.45 | 1.17 (1.68 - 0.70) |
| CAT + Pois. (autoc.) | 3.8 | 2.70 | 1.12 (1.50 - 0.75) |
| CAT + Pois. (uncor.<sup>b</sup>) | 3.5 | 2.45 | 1.94 (2.62 - 1.26) |
| CAT + Pois. (uncor.) | 3.5 | 2.70 | 1.84 (2.46 - 1.22) |
| CAT + Pois. (uncor.) | 3.8 | 2.45 | 1.67 (2.31 - 1.03) |
| CAT + Pois. (uncor.) | 3.8 | 2.70 | 2.03 (2.70 - 1.38) |

<sup>a</sup>Log normal autocorrelated clock model.<sup>b</sup>Uncorrelated gamma model

To understand the effects of the oldest calibration point (point 11, Fig. 3) on the estimated divergence time, we tested a second set of boundaries restricting this point to a minimum age of 2.7 Ga (Calibration 2) to consider the possibility that the record for oxygen several hundred million years before the GOE was produced by crown group Cyanobacteria (Planavsky et al., 2014; Havig et al., 2017). Supplementary Fig. S2 provides a comparison of the two calibration choices on the overall estimated times for flexible and non-flexible models. If the divergence times using both calibrations are plotted against each other, a linear relationship is obtained (see supplementary Table S6 for linear regression). Calibration 2 did not seem to have a very strong effect on the estimated divergence times nor **ΔT**. For example, under a root prior of 3.5 ± 0.05 Ga and employing Calibration 2, **ΔT** was centered at 0.97 Ga (Table 4). We also tested the effect of removing the oldest calibration point from the analysis (point 11, Fig. 3), this has the effect of shifting of making many nodes younger, yet **ΔT** remained in the range of 0.8 to 1.3 Ga depending on the level of flexibility allowed (Table 1).

In contrast, the choice of model for the evolution of substitution rates had a strong impact on the estimated divergence times as shown in supplementary Fig. S3 and Table S7. Supplementary Fig. S3 presents a comparison of divergence time estimates of a tree calculated using a relaxed log-normal autocorrelated molecular clock with a tree calculated using an uncorrelated gamma model on the rates of evolution. The autocorrelated model assumes that neighboring branches are more likely to evolve at a similar rate, while the uncorrelated model assumes that the rate of evolution of each branch can vary independently (Lepage et al., 2007; S. Y. W. Ho & Duchene, 2014). Under the uncorrelated model the estimated divergence times of many nodes were aberrantly shifted to younger ages: for example, most Cyanobacterial and eukaryotic D1 clustered in the range of 700 to 0 Ma, which is inconsistent with the fossil record. The molecular mechanism behind this difference could be related to the fact that photochemistry imposes a strong constraint on the evolution of reaction center proteins: as all of them must coordinate and maintain all redox cofactors, chlorophylls, quinones, carotenoids, and a non-heme iron, at a precise orientation and distance from each other to allow for control of electron transfer rates and redox potentials. These rates and potentials are crucial not only for function but also for protection against the formation of reactive oxygen species (Cardona et al., 2012; Rutherford et al., 2012). It seems reasonable then, that the rates of evolution of all Type II reaction center proteins should be similar between closely related groups, thus corresponding to an autocorrelated model.

## Discussion

### Change in sequence identity as a function of time

As an approximation to the evolution of the core subunits of PSII we plotted the level of sequence identity of D1 and D2 as a function of known divergence times (Fig. 2). Two main conclusions can be derived from this plot, which are also in agreement with the molecular clock analysis. Firstly, the three earliest stages in the evolution of oxygenic photosynthesis: the divergence of Type I and Type II reaction centers, the divergence of *anoxygenic* and *oxygenic* families of Type II reaction center proteins, and the divergence of D1 and D2, are more likely to have started soon after the origin of the first reaction centers rather than near the GOE. Taking into consideration that there is strong evidence for photosynthesis at 3.5 Ga (Tice & Lowe, 2004; Nisbet & Fowler, 2014; Butterfield, 2015; Schopf et al., 2018), then all three stages could well predate this time. Secondly, the MRCA of Cyanobacteria is more likely to have lived near the time of the GOE rather than shortly after the origin of photosynthesis in the early Archean. This common ancestor must have had a standard PSII, which places it relatively far after the origin of photosynthesis. The early divergence of D1 and D2 means that the earliest stages in the evolution of oxygenic photosynthesis could predates the MRCA of Cyanobacteria by over a billion years.

### Bayesian relaxed molecular clock analysis

The application of a Bayesian molecular clock analysis to the phylogeny of Type II reaction centers is strongly constrained by three pieces of well-supported evidence: 1) evidence of photosynthesis at 3.5 Ga, 2) by strong evidence for the GOE at 2.4 Ga, and 3) by the relatively slow rate of evolution of the core proteins over the Proterozoic. Under these constraints, the divergence between D1 and D2 is better explained by the duplication event occurring early in the evolutionary history of photosynthesis, in the early Archean, with the appearance of standard PSII occurring after a long period of evolutionary innovation. We highlight this long period by introducing the concept of **ΔT**. The magnitude of **ΔT** is dictated by the large phylogenetic distance between D1 and D2 and the relatively slow rate of evolution determined from the geochemical and fossil record. Therefore, it is not surprising that under most models employed in this analysis **ΔT** is in the range of 1.0 Ga.

We have considered two possible evolutionary scenarios that are both consistent with a large **ΔT** (Fig. 6). In the first scenario the standard forms of D1 start to diverge at about 2.4 Ga, as seen in Fig. 3, and diversify into G3 and G4 after the GOE. If we consider that the MRCA of Cyanobacteria had a G4 D1, this would set it after the GOE. This scenario, derived from the application of a relaxed molecular clock using a non-parametric CAT model with flexible boundaries, is in agreement with the recent observations by Shih, Hemp, et al. (2017) and other molecular clock studies that placed the divergence of *Gloeobacter* after the GOE (Feng et al., 1997; David & Alm, 2011; Marin et al., 2017). In this scenario, assuming that the earliest events in the history of photosynthesis started about 3.5 Ga, the divergence of D1 and D2 is set at about 3.2 Ga.

**Figure 6.**
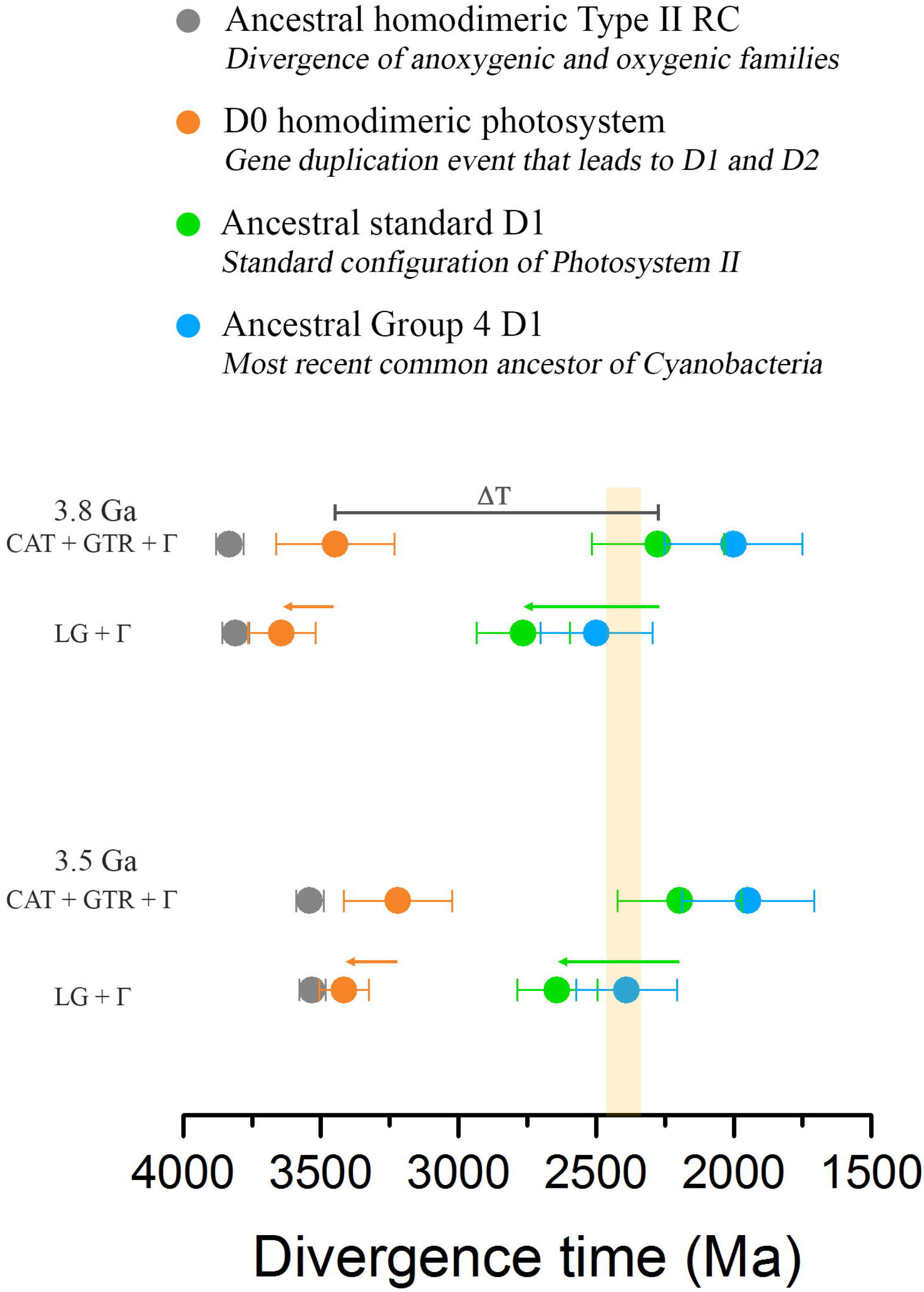
Scenarios for the evolution of Type II reaction centers. The rows of colored dots represent estimated divergence times of key nodes as highlighted in Fig. 3 and calculated using the CAT + GTR + Γ and LG + Γ models and root priors of 3.8 or 3.5 ± 0.05 Ga. A highly oxidizing photosystem with enough power to split water is likely to have originated before the gene duplication event that led to D1 and D2 (orange dot). Making the MRCA of Cyanobacteria older (green arrow) pushes the earliest stages in the evolution of PSII and water oxidation closer to the origin of photosynthesis (orange arrow). The yellow vertical bar marks the GOE

In the second scenario, we considered that the MRCA of Cyanobacteria occurs before the GOE as suggested by other molecular clock analyses (Falcon et al., 2010; Sanchez-Baracaldo, 2015; Schirrmeister et al., 2015). In the present work, this scenario can be fitted most satisfactorily with the application of a relaxed molecular clock using an empirical amino acid substitution model (LG + Γ).

In this scenario, under a root prior of 3.5 Ga, the appearance of the ancestral standard form of D1 is set at about 2.6 Ga; and this has the consequence of pushing the divergence of D1 and D2 closer to the root, and thus **D0** is set at about 3.4 Ga (Table 1 and 2, Fig. 6). This effect is due in part to the fact that the phylogenetic distance between D1 and D2 is invariable, and thus under any scenario the data is better explained by a long span of time separating **D0** and the standard heterodimeric PSII. What can be concluded from this is that the older the MRCA of Cyanobacteria is, the more likely it is that the divergence of D1 and D2 started near the origin of photochemical reaction centers and thus, near the origin of photosynthesis.

### Rates of evolution

It should be observed that if a relatively constant rate of evolution were to be applied to the phylogeny of Type II reaction centers, the root of the tree would be placed long before the formation of the planet (dashed red line in Fig. 2). Given the fact that the rate of evolution of D1 and D2 is constrained at comparatively slow rates from the Proterozoic until present times, the period of fast evolutionary change has to be located at the earliest stages to account for the evolution of photosynthesis within a reasonable timeframe. We found that even considering an origin of Type II reaction centers at 3.8 Ga the maximum rate during the evolution of D1 and D2, **ν**_**max**_, is more than thirty times greater than the measured rates since the Proterozoic.

The phenomenon of an initial fast rate of evolution followed by an exponential decrease demands an explanation. A mechanism that can lead to accelerated rates of evolution is gene duplication followed by functional innovation (Lynch & Conery, 2000; Innan & Kondrashov, 2010). In such scenario the fast rates of evolution at the nodes representing the ancestral duplication of **K** and **D0** may be linked to the early origin of the very first Type II reaction centers and the initiation of heterodimerization in each respective case. Studies of the acceleration of the rates of evolution suggests that immediately after a gene duplication event there is an acceleration of the gene copy that acquires a new function followed by a slow-down of the rates (Duda & Palumbi, 1999; Lynch & Conery, 2000; Rosello & Kondrashov, 2014): however, the timing of these events seem to be completed within tens of millions of years (Rosello & Kondrashov, 2014) and not in the scale of a billion years as detected for the evolution of the core Type II reaction center proteins (Fig. 5 A). Nevertheless, it is plausible that major evolutionary innovations such as the origin of photosynthesis or aerobic respiration may have had more pronounced and prolonged effects in the rate of evolution of duplicated genes essential for these processes.

Another possibility is the temperature dependent deamination of cytosine, as suggested by Lewis and coworkers (2016). They calculated that as the Earth cooled during its first 4 Ga, the rate of spontaneous mutation would have fallen exponentially by a factor of more than 4000. That is to say that the rate of spontaneous mutation during the earliest stages in the history of life would have been about three orders of magnitude greater than those observed since the Proterozoic. Lewis and coworkers (2016) calculated that 50% of all spontaneous mutations occurred in the first 0.2 Ga, which matches well with the exponential decay trend seen in Fig. 5A, especially if an origin of photosynthesis is considered to be about 3.8 Ga. A third possibility is higher UV radiation on the planet’s surface during the early Earth in the absence of an ozone layer, which could have resulted in rates of DNA damage up to three orders of magnitude greater than in present day Earth, as calculated by Cockell (2000); this higher rate of damage may have led as well to faster rates of change.

### Diversification of phototrophic lineages

Cardona (2015) suggested based on the phylogeny of reaction center proteins that some of the earliest events in the evolution of photosynthesis, like the divergence of Type I and Type II reaction centers, predated the diversification event that gave rise to the known diversity of phototrophs. In accord with this, several recent studies have highlighted that many, if not all, the described groups of phototrophs appear to be much younger than the origin of photosynthesis itself, with every group having close non-photosynthetic relatives (Fischer et al., 2016). For example, Shih, Hemp, et al. (2017) reported a molecular clock of Cyanobacteria and their closest non-photosynthetic relatives the Sericytochromatia and the Melainabacteria (Soo et al., 2014; Soo et al., 2017), in which the divergence of *Gloeobacter* was estimated to postdate the GOE, at about 2.0 Ga. Such late dates seem to contrast with other recent molecular clock studies that placed the same node before the GOE, in the range of 2.8 to 2.6 Ga (Sanchez-Baracaldo, 2015). Along the same lines, Magnabosco et al. (2018) estimated that the divergence of *Gloeobacter* likely occurred around the GOE, between 2.7 to 2.2 Ga depending on the calibration choice and evolutionary model. A different molecular clock study on the evolution of the phylum Chloroflexi suggested the phototrophy arrived to this phylum via HGT about 0.9 Ga ago (Shih, Ward, et al., 2017). Consistent with this, Magnabosco et al. (2018) estimated that phototrophic Chloroflexi diverged between 2.4 and 1.0 Ga, depending on the models used. In the same study, Magnabosco et al. (2018) also timed the origin of phototrophic Chlorobi, having homodimeric Type I reaction centers, at about 1.7 Ga, which is consistent with biomarker evidence (Brocks et al., 2005). It is worth noting, however, that even accounting for high levels of uncertainty it looks like the most recent common ancestors of each of these phototrophic groups, anoxygenic or oxygenic, occurred long after the origin of photosynthesis.

The age estimates derived from the evolution of Type II reaction centers presented here are consistent with the molecular clock studies mentioned above and at the same time emphasize the antiquity of photosynthesis. More specifically, our data suggests that the standard form of PSII core proteins, inherited by all photosynthetic Cyanobacteria, started to diversify around the GOE. Moreover, and as shown in Fig. 3, our timing for the start of the diversification of Chloroflexi-type L and M subunits is placed after the GOE. In an independent study, a molecular clock analysis of Type I reaction center proteins placed the MRCA of phototrophic Chlorobi after the GOE, in the range of 2.1 and 1.6 Ga (Cardona, 2018).

The data presented in this work adds an extra dimension to the timing of the emergence of the known described groups of phototrophs by following the history of the reaction centers themselves as direct proxies for the evolution of photosynthesis. We can conclude then that when each of the known groups of phototrophs diversified, and this includes Cyanobacteria as well, their reaction centers had already reached a high-degree of specialization regardless of any gains or losses of phototrophy along this evolutionary trajectory. From this perspective, if the duplication of **D0** occurred in the early Archean it is possible that the Sericytochromatia and Melainbacteria classes of non-photosynthetic Cyanobacteria originated after losses of oxygenic photosynthesis, as they specialized in heterotrophic lifestyles and were supported by the primary productivity of their photosynthetic cousins. The loss of oxygenic photosynthesis is a relatively common process across the tree of life, several well-known examples are: the secondary loss of all PSII genes by the symbiont *Atelocyanobacterium talassa* (Thompson et al., 2012; Cornejo-Castillo et al., 2016), the complete loss of photosynthesis of the cyanobacerium endosymbiont of *Epithemia turgida* (Nakayama et al., 2014), the case of parasitic Apicomplexa (Moore et al., 2008) and holoparasitic angiosperms (Ravin et al., 2016).

### Was there water oxidation before D1 and D2?

Before the gene duplication that allowed the divergence of D1 and D2, the ancestral homodimeric photosystem had enough oxidizing power to form the neutral tyrosyl radical. High enough to surpass the E_*m*_ of water oxidation to oxygen, but this does not necessarily imply that water oxidation was occurring at this time. Is there any evidence that would support or disprove an origin of water oxidation before the D1 and D2 duplication event?

Almost every major structural difference between anoxygenic Type II reaction center proteins and the core proteins of PSII can be explained in the context of water oxidation, protection against the formation of reactive oxygen species, and enhanced repair and assembly mechanisms due to oxidative damage from the formation of singlet oxygen around the photochemical pigments (see below). Five major structural differences distinguish D1 and D2 from L and M (Fig. 7 A and B). Starting from the N-terminus:

**Figure 7.**
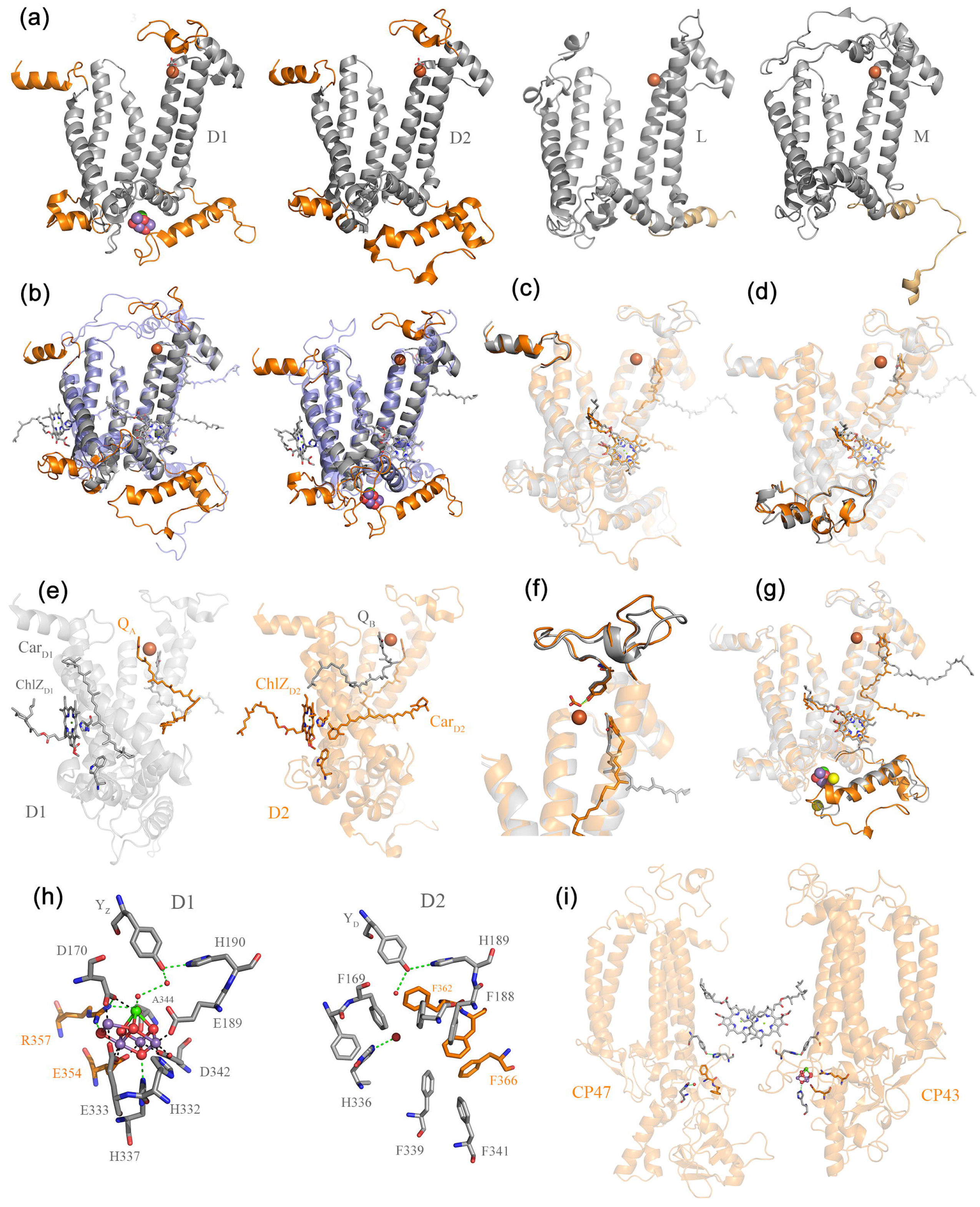
Structural comparisons of Type II reaction center proteins. (*a*) Several structural domains are conserved in D1 and D2 but are absent in L and M: these are highlighted in orange. D1 and D2 are plotted from the crystal structure of PSII from *Thermosynechococcus vulcanus*, PDB: 3WU2 (Umena et al., 2011) and L and M from *Thermochromatium tepidum*, PDB: 3WMM (Niwa et al., 2014). (*b*) Overlap of D2 (gray) with M (transparent blue) and D1 (gray) with L (transparent blue). (*c*) Overlap of D1 (gray) and D2 (orange) highlighting the conserved N-terminus. (*d*) Overlap of D1 and D2 highlighting the conserved protein fold between the 1^st^ and 2^nd^ transmembrane helices. (*e*) D1 is shown in gray and D2 in orange. ChlZ_D1_ Car_D1_ W105, and Q_B_ are shown in gray sticks. ChlZ_D2_, Car_D2_, W104, and Q_A_ are shown in orange sticks. (*f*) Overlap of D1 and D2 highlighting the conserved protein fold where the bicarbonate binding site is placed. (*g*) Overlap of D1 and D2 highlighting the conserved protein fold at the C-terminus. (*h*) The Mn_4_CaO_5_ cluster coordination sphere and the equivalent location in D2. (*i*) Perspective view showing the interaction of CP47 and CP43 with the electron donor side of D2 and D1 respectively. The reason why CP47, and in particular CP43, interact with the donor side of PSII is an unsolved mystery given the fact that their main role is that of light harvesting. It can be rationalized however if water oxidation started in a homodimeric reaction center early during the evolution of photosynthesis (Cardona, 2017)

*The N-terminus itself* (Fig. 7 C). PSII is known to generate singlet oxygen, a very damaging form of reactive oxygen species that without control can lead to irreparable damage to the organism and death. Singlet oxygen is produced when molecular oxygen interacts with the excited triplet state of chlorophyll (Krieger-Liszkay et al., 2008; Vass & Cser, 2009; Rutherford et al., 2012). Triplet chlorophyll is in turn formed when excess harvested light energy cannot be efficiently dissipated or when the forward electron transfer reactions of PSII are blocked and instead a backflow of electrons occurs (back-reactions) (Santabarbara et al., 2002; Krieger-Liszkay et al., 2008). Thus, the unavoidable production of singlet oxygen by PSII results in rapid damage of the core proteins in such a way that the half-lifetime of D1 is about 30 minutes. D1 is known to be the protein with the fastest turnover in the photosynthetic membrane (Aro et al., 1993). The half-lifetime of D2 is also relatively fast measured at about 3 hours. In comparison, the half-lifetime of Photosystem I core proteins is about 2 days (Yao et al., 2012). Damaged D1 and D2 are degraded by dedicated FtsH proteases, which target and recognize the N-terminus of both subunits. Deletion of the N-terminus results in impairment of degradation and repair (Komenda et al., 2007; Krynicka et al., 2015). The preserved sequence and structural identity at the N-terminus of both D1 and D2 suggests that the evolution of enhanced repair mechanisms had started to evolve before the duplication. Consistent with this, the evolution of all bacterial FtsH proteases confirms that the lineage of proteases specifically dedicated to the repair of PSII make a monophyletic and deep-branching clade (Shao et al., 2018). As it is the case for the evolution of reaction center proteins, this deep-branching clade of PSII-FtsH proteases appeared to have diverged before the radiation of those found in all groups of phototrophs (Shao et al., 2018).

*A protein fold between the 1^st^ and 2^nd^ transmembrane helices* (Fig. 7 D). In D1 and D2 this region is made of 54 and 52 residues, respectively, in comparison to 28 and 35 residues in L and M, respectively. This fold is enlarged in D1 and D2 to provide a site for protein-protein interactions with the small peripheral subunits and the extrinsic polypeptides (Cardona, 2015, 2016), none of which are present in anoxygenic Type II reaction centers. In D1 this site provides a connection to PsbI, M, T and O; and in D2 to the Cytochrome *b*_*559*_, PsbH, J and X. The small subunits are necessary to support a more complex assembly and disassembly cycle due to much higher rates of repair (Komenda et al., 2012). They provide stability, help with photoprotective functions, assist with the photoassembly of the Mn_4_CaO_5_ cluster (Komenda et al., 2002; Dobakova et al., 2007; Popelkova & Yocum, 2011; Hamilton et al., 2014; Sugiura et al., 2015), and even contribute to the highly oxidizing potential of PSII (Ishikita et al., 2006). The extrinsic polypeptides are fundamental for the stability of the Mn_4_CaO_5_ cluster, in particular PsbO, also known as the manganese stabilizing protein (Franzen et al., 1985; De Las Rivas et al., 2004; Roose et al., 2016). That this site and its structural fold is conserved in D1 and D2 suggests that before their divergence the ancestral homodimeric photosystem had already achieved a high degree of structural complexity and was interacting with a number of auxiliary subunits in a way that it is not matched by anoxygenic Type II reaction centers. Because the role of the auxiliary subunits is the support of water oxidation and associated functions, it seems likely that this expansion of structural complexity would only start after the origin of water oxidation.

*The peripheral pigment pairs, ChlZ_D1_-Car_D1_ and ChlZ_D2_-Car_D2_* (Fig. 7 E). D1 and D2 each coordinate a peripheral chlorophyll from a conserved histidine ligand in the 2^nd^ transmembrane helix, known as ChlZ_D1_ and ChlZ_D2_. These peripheral pigments are absent in anoxygenic Type II reaction centers but are present in Type I reaction centers indicating that they existed before the divergence of D1 and D2 (Cardona, 2015). Both peripheral chlorophylls are required for photoautotrophic growth as mutations that impair their binding cannot assemble functional PSII (Lince & Vermaas, 1998; Ruffle et al., 2001). ChlZ_D1_ and ChlZ_D2_ are each in direct contact with a beta-carotene molecule, known as Car_D1_ and Car_D2_ respectively, seen using crystallography first by Ferreira et al. (2004) and Loll et al. (2005), but detected and characterized by spectroscopy well before that, see for example (Kwa et al., 1992; Noguchi et al., 1994; Hanley et al., 1999). The position of Car_D1_ and Car_D2_ differs in that the former is positioned perpendicular to the membrane plane while the latter is parallel to the membrane plane: however, one of the beta-rings of each carotenoid links to ChlZ_D1_ and ChlZ_D2_ via strictly conserved tryptophan residues (D1-W105 and D2-W104, respectively), located in the unique protein fold between the 1^st^ and 2^nd^ transmembrane helices described above, and therefore absent in L and M. Car_D2_ is tilted relative to CarD1 partly to give way to the exchangeable plastoquinone, Q_B_. The core carotenoids of PSII have been shown to contribute little to light-harvesting and have dominantly a protective role (Stamatakis et al., 2014). The close association of ChlZ_D1_ and ChlZ_D2_ with carotenoids would suggest a role in protection, quenching chlorophyll triplet states or directly scavenging singlet oxygen (Cogdell et al., 2000; Telfer, 2002). A role for direct quenching of singlet oxygen for both ChlZ_D1_-Car_D1_ and ChlZ_D2_-Car_D2_ has been suggested based on spectroscopy of isolated reaction centers (Telfer et al., 1994). Furthermore, ChlZ_D2_-Car_D2_ have been demonstrated to be involved in protective electron transfer side pathways within PSII. For a detailed overview of these pathways see for example (Faller et al., 2005). That ChlZ_D1_ and ChlZ_D2_ have been retained since before the divergence of Type I and Type II reaction centers indicates that they predate the D1 and D2 divergence. The acquisition of closely interacting carotenoids seem to have occurred therefore after the **K** and **D0** divergence, but before the D1 and D2 split, in support of water oxidation before heterodimerization.

*An extended loop between the 4^th^ and the 5^th^ transmembrane helices* (Fig. 7 F). This is required for the coordination of bicarbonate, a ligand to the non-heme iron (Ferreira et al., 2004), which is a distinctive feature of PSII. In anoxygenic Type II reaction centers the non-heme iron is coordinated instead by a glutamate from the M subunit. There is significant sequence and structural conservation of the bicarbonate binding site in D1 and D2. Two strictly conserved tyrosine residues D1-Y246 and D2-Y244 provide hydrogen bonds to bicarbonate (Ferreira et al., 2004). This indicates that bicarbonate binding was a feature existing before the divergence of D1 and D2. The role of bicarbonate had been a long-standing mystery, but recently it was shown that binding and unbinding of bicarbonate modulates the E_*m*_ of the quinones, working as a switch from a productive state into a protective state that prevents chlorophyll triplet state and singlet oxygen formation (Brinkert et al., 2016). Previously, G. N. Johnson et al. (1995) had shown that a shift in the E_*m*_ of the fixed quinone, Q_A_, plays a key role in protection of PSII during assembly of the Mn_4_CaO_5_ cluster, a light-driven process. It is understood now that such a shift is mediated by bicarbonate binding (Brinkert et al., 2016). A further conclusion from this is that the ancestral photosystem made of a **D0** homodimer had already evolved to incorporate bicarbonate-mediated protective mechanisms as well, implying oxygen evolution, and by extension, the assembly of a primordial water-oxidizing complex.

*An extended C-terminus and the Mn_4_CaO_5_ cluster binding site* (Fig. 7 G). D1 and D2 share an extended C-terminus made of about 50 residues and with a distinctive alpha-helix parallel to the membrane plane. The C-terminus is necessary for the coordination of the Mn_4_CaO_5_ cluster, Cl^−^ binding, water channels, and proton pathways (Debus, 2001; Umena et al., 2011; Linke & Ho, 2013). Remnants of this C-terminal extension may be found in some of the M and L subunits of phototrophic Proteobacteria and Chloroflexi (Cardona, 2015), but see Fig. 7 A. In D1 H332, E333, D342 and A344 coordinate the Mn_4_CaO_5_ cluster (Fig. 7 H). H337 provide a hydrogen bond to one of the bridging oxygens. In addition, E354 from the CP43 antenna subunit coordinates two of the Mn atoms and R357 offers a hydrogen bond to another bridging oxygen. While there is not a cluster in D2, an examination of the donor side in the immediate vicinity of the redox active tyrosine shows that the site has been blocked by a number of phenylalanine residues. Every ligand to the cluster is matched by a phenylalanine residue in D2 (Fig. 7 H). In the CP47 antenna the ligands found in the CP43 are also replaced by phenylalanine residues (Fig. 7 H and I). The only exception is a histidine in a position equivalent to H337, which perhaps not coincidentally, provides a hydrogen bond to a water molecule locked within the hydrophobic patch made by the phenylalanine residues. No such phenylalanine patch is found in any other reaction center protein, except D2 (Cardona, 2016). The presence of a redox tyrosine in D2 and what seems like a vestigial metal-binding site would be puzzling if the water-oxidizing cluster evolved after the divergence of D1 and D2 in a heterodimeric system, but it would make sense if a primordial water-oxidizing cluster appeared first on both sides of the reaction center in the ancestral homodimeric photosystem.

Based on the above structural and functional evidence we find it likely that water oxidation originated before the divergence of D1 and D2. Hence, in a homodimeric Type II ancestor containing two identical exchangeable quinones charge separation would result in enhanced back-reactions caused by electron transfer to a quinone site when empty or when occupied by a reduced form of the quinone; and additionally, by shorter back-reaction pathways (Cardona et al., 2012; Rutherford et al., 2012). Back-reactions would give rise to chlorophyll triplet states in the heart of the reaction center (Dutton et al., 1972; Rutherford et al., 1981). In the evolution of a water-splitting homodimeric ancestor with an increased oxidizing potential, as mentioned above, avoidance of photodamage could be a significant selective pressure for heterodimerization, as the enhanced back-reactions intrinsic to the homodimeric Type II reaction centers would have become a major problem whenever oxygen was present. An inefficient homodimeric water-splitting photosystem, would have encountered this problem first and thus come under strong selection pressure toward heterodimerization at an early time. The results of the present work fit with this picture. Furthermore, our present finding that heterodimerization started significantly later in anoxygenic Type II reaction centers, can be explained in this context as well, since **K** would have encountered these selection pressures much later, at a time when oxygen concentrations began to rise in localized environments in the late Archean period, nearing the GOE (Bosak et al., 2009; Lyons et al., 2014; Riding et al., 2014).

If water oxidation started in a homodimeric photosystem, this would place the earliest stages in the evolution of oxygenic photosynthesis in the early Archean more than a billion years before the GOE. It follows then that the heterodimerization process of D1 and D2 was likely a path to optimization of water oxidation efficiency (Rutherford et al., 2012), photoprotective mechanisms (Krieger-Liszkay et al., 2008), photoactivation of the Mn_4_CaO_5_ cluster, assembly, and repair of the protein complex in the presence of oxygen (Nixon et al., 2010; Nickelsen & Rengstl, 2013).

## Final remarks

The evolution of Type II reaction centers highlights the long history of oxygenic photosynthesis before the Great Oxidation Event. We showed that the span of time between the gene duplication event that led to D1 and D2 and the appearance of standard PSII could have been about a billion years. We argued that water oxidation is likely to have started before the divergence of D1 and D2. So what happened during this long period of time? If water oxidation originated in a simpler homodimeric photosystem in a completely anaerobic world, the large increase in the structural complexity of PSII, PSI, and associated light harvesting complexes had to occur along this trajectory. This includes the acquisition of many peripheral and extrinsic protein subunits and the heterodimerization of D1 and D2, CP43 and CP47, and PsaA and PsaB. At the same time, the thermodynamic coupling between both photosystems and the retuning of the entire electron transport chain and all electron carriers to increasingly oxidizing conditions also had to occur. This expansion in complexity had to be coupled with the evolution of highly organized assembly and repair processes. Thus, the first water-oxidizing reaction centers may have been active only for brief amounts of time in the absence of efficient repair. Greater water oxidation efficiency also needed the innovation of photoprotective mechanisms acting at different time scales spanning several orders of magnitude, like dissipatory recombination pathways, non-photochemical quenching, or state-transitions. Furthermore, the light reactions of photosynthesis had to be linked to carbon fixation and other downstream metabolic process. Signaling, feedback, and regulatory mechanisms had to be put in place to control photosynthesis under varying environmental conditions and across a diel cycle. Needless to say, all anaerobic reactions and processes inhibited by oxygen originally found in the earliest anaerobic water-oxidizing ancestors had to be separated from oxygen production or readapted to work under aerobic conditions. The link to carbon fixation is of particular importance. CO_2_ levels in the atmosphere were higher than now (Sheldon, 2006; Nutman et al., 2017). However, limiting diffusion across boundary layers in mats and stromatolites would have restricted anoxygenic and early oxygenic phototrophs alike. The development of water oxidation would have opened up the way to faster photosynthetic rates, spurring on gross primary production rates later in the Archean, with the concomitant need for increases in nitrogen fixation.

## Materials and Methods

### Phylogeny of Type II reaction centers

Sequences were retrieved from the RefSeq NCBI database using PSI-BLAST restricted to Cyanobacteria, Proteobacteria, and Gemmatimonadetes. A total of 1703 complete sequences were downloaded and aligned using Clustal Omega employing ten combined guide trees and Hidden Markov Model iterations (Sievers et al., 2011). To confirm that the alignment conformed with known structures, the 3D structures of the D1, D2, L and M, from the crystal structures 3WU2 (Umena et al., 2011) of PSII and 2PRC (Lancaster & Michel, 1997) of the anoxyenic Type II reaction center were overlapped using the CEalign (Jia et al., 2004) plug-in for PyMOL (Molecular Graphics System, Version 1.5.0.4 Schrödinger, LLC) and structural homologous positions were cross-checked with the alignment. Maximum Likelihood (ML) phylogenetic analysis was performed using PhyML 3.1 (Guindon et al., 2010) using the LG model of amino acid substitution. The amino acid equilibrium frequencies, proportion of invariable sites, and across site rate variation were allowed to be determined by the software from the dataset. Tree search operations were set as the best from the Nearest Neighbor Interchange and Subtree Pruning and Regrafting method, with the tree topology optimized using five random starts. The ML tree using all sequences is shown in Fig. 1 and it replicates earlier evolutionary studies of Type II reaction centers that used simpler methods and fewer sequences (Beanland, 1990; Cardona, 2015).

### Change in sequence identity as a function of time

To get a better understanding of the evolutionary trends of D1 and D2 reaction center proteins as a function of time, we compared the percentage of sequence identity of D1 and D2 proteins from species of photosynthetic eukaryotes with known or approximate divergence times. In total 23 pairs of sequences were compared and are listed in Supplementary Table 1. When two sequences were of different length, the longest was taken as 100%. Of these 23 pairs, the first 16 were based on the fossil calibrations recommended by Clarke et al. (2011) after their extensive review of the plant fossil record. Divergence times were taken as the average of the hard minimum and soft maximum fossil ages suggested by the authors. The last 7 comparisons are approximate dates taken from a recent molecular clock analysis of the evolution of red algae E. C. Yang et al. (2016). The plot of sequence identity *vs* approximate divergence time was then fitted with a linear regression and the fitting parameters are shown in supplementary Table S2.

### Bayesian relaxed molecular clock and fossil calibrations

A total of 54 bacterial sequences spanning the entire diversity of Type II reaction centers were selected for Bayesian molecular clock analysis, including atypical and standard forms of D1, Alpha-, Beta-, Gammaproteobacteria, Chloroflexales, and *Gemmatimonas phototrophica*. Furthermore, 10 D1 and 10 D2 sequences from photosynthetic eukaryotes from taxa with a well characterized fossil record were added to allocate calibration points. The phylogeny of Type II reaction centers was cross-calibrated on D1 and D2 as listed in Table 5 and calibration points were assigned as presented in Fig. 3, red dots.

**Table 5.** Calibration points

| **Node** | **Event** | **Maximum (Ma)** | **Minimum (Ma)** |
| --- | --- | --- | --- |
| **1** | *Arabidopsis*-*Populus* divergence | 127 | 82 |
| **2** | Angiosperms | 248 | 124 |
| **3** | Gymnosperms | 366 | 306 |
| **4** | Land plants | - | 475 |
| **5** | Diatoms | - | 190 |
| **6** | Floridæ | - | 600 |
| **7** | MRCA of red algae | - | 1200 |
| **8** | Heterocystous Cyanobacteria | - | 1600 |
| **9** | Pleurocapsales | - | 1700 |
| **10** | Early-branching multicellular Cyanobacteria | - | 1900 |
| **11** | MRCA of Cyanobacteria | - | 2450/2700 |

Dates for the *Arabidopsis/Populus* divergence, the divergence of the angiosperms (*Amborella*), gymnosperms (*Cycas*), and land plants (*Marchantia*) were implemented as suggested and discussed by Clarke et al. (2011), representing points 1 to 4 in Fig. 3. Three ages from eukaryotic algae were used too: an age of 190 Ma was assigned to the divergence of *Phaeodactylum trichornutum* and *Thalassiosira pseudonana*, based on fossil Jurassic diatoms from the Lias Group, reviewed by Sims et al. (2006). A minimum age of 600 Ma based on a Late Neoproterozoic Chinese multicellular red alga fossil (Xiao et al., 2004) was assigned to the split between the diatom sequences and the sequences from *Porphyra purpurea*, as a conservative estimate for the divergence of complex red algae, which predates this time (E. C. Yang et al., 2016). The oldest calibration point in photosynthetic eukaryotes was assigned as a minimum age of 1.2 Ga to the divergence of the sequences from *Cyanidium caldarium* a unicellular early-branching red algae. This was used as a conservative estimate for the origin of photosynthetic eukaryotes. The earliest widely accepted fossil of a photosynthetic eukaryote is that from a multicellular red algae, *Bangiomorpha* (Butterfield, 2000; Knoll et al., 2013), thought to be 1.0 Ga (Gibson et al., 2017). Recently described multicellular eukaryotic algae fossils have been reported at 1.6 Ga (Bengtson et al., 2017; Qu et al., 2018; Sallstedt et al., 2018) suggesting that the earliest photosynthetic eukaryotes might be older than that, which would be consistent with recent molecular clock analysis (E. C. Yang et al., 2016; Sánchez-Baracaldo et al., 2017).

Previously implemented Cyanobacterial fossils were also used to calibrate the clock (Blank & Sanchez-Baracaldo, 2010; Sanchez-Baracaldo, 2015; Sánchez-Baracaldo et al., 2017). A minimum age of 1.6 Ga was assigned to Nostocales because described akinetes of this age have been found in cherts from Siberia, China, and Australia (Golubic et al., 1995; Tomitani et al., 2006; Schirrmeister et al., 2016). Pleurocapsales are characterized by multiple fissions during cell division and fossils retaining this morphology have been described at 1.7 Ga (Golubic & Lee, 1999; Sergeev et al., 2002). The earliest well-assigned filamentous Cyanobacteria fossils of comparable size to those of the early-branching *Pseudanabaena* have been reported at 1.9 Ga (Golubic & Lee, 1999; Sergeev et al., 2002; Schirrmeister et al., 2016; Sánchez-Baracaldo et al., 2017), and this was assigned as a minimum age to the sequences from *Pseudanabaena biceps*.

The oldest calibration point, point 11, was selected to be the branching point of the D2 and the G4 D1 from *Gloeobacter violaceous.* This was set to be around the age for the GOE and was assigned as a minimum age of 2.45 Ga (Calibration 1) (Bekker et al., 2004). For comparison, a calibration of 2.7 Ga was also used (Calibration 2) to test the effect on the estimated divergence times of an older age for crown group Cyanobacteria. Geological evidence suggests that oxygen ‘whiffs’ or ‘oases’ could significantly predating the GOE (Lyons et al., 2014; Planavsky et al., 2014; Havig et al., 2017) so this scenario is not entirely implausible.

The calibration points on D1 were allocated on Group 4 because this type of D1 is the only one retained in all Cyanobacteria with PSII, it is the only type of D1 inherited by photosynthetic eukaryotes, and it is the main D1 used to catalyze water oxidation under most growth conditions.

It is well accepted that a form of anoxygenic photosynthesis had already evolved by 3.5 Ga. This is demonstrated by both sedimentological and isotopic evidence for photoautotrophic microbial communities recorded in Paleoarchean rocks (Tice & Lowe, 2004; Nisbet & Fowler, 2014; Butterfield, 2015). In addition, sedimentary rocks and banded iron formations from Isua, Greenland, hint at the presence of photosynthetic bacteria in the marine photic zone as early as 3.7-3.8 Ga (Schidlowski, 1988; Rosing, 1999; Rosing & Frei, 2004; Grassineau et al., 2006; Czaja et al., 2013; Knoll, 2015). Therefore, we used a root prior of 3.5 Ga as a conservative estimate for photoautrophy based on photochemical reaction centers. Nevertheless, because it is not yet known exactly when photosynthesis originated for the first time we also tested the effect of varying the root prior from 3.2 to 4.1 Ga on the estimated divergence time under restrictive and flexible scenarios.

A Bayesian Markov chain Monte Carlo approach was used to calculate the estimated divergence times. We used Phylobayes 3.3f, which allows for the application of a relaxed log-normal autocorrelated molecular clock under the CAT + GTR + Γ model (Lartillot & Philippe, 2004; Lartillot et al., 2009) necessary for the implementation of flexible boundaries on the calibration points (Z. Yang & Rannala, 2006). To understand the effect of different evolutionary models on the age estimates we compared the CAT + GTR + Γ model with 1) a LG + Γ model that sets less flexible boundaries on the calibration points, 2) a CAT + Γ model assuming a uniform (Poisson) distribution of amino acid equilibrium frequencies, or 3) an uncorrelated gamma model where the rates of substitution can vary independently. The flexible bounds on the CAT + GTR + Γ model were set to allow for 2.5% tail probability falling outside each calibration boundary or 5% in the case of a single minimum boundary. All molecular clocks were computed using four discrete categories for the gamma distribution and four chains were run in parallel until convergence.

In this work we define the period of time between the duplication event that led to the divergence of D1 and D2 and the appearance of the ancestral standard D1 as **ΔT**. This value is calculated as the subtraction of the mean age of the latter node (Fig. 3, green dot) from the former’s mean node age (Fig. 3, **D0**, orange dot). The instant rates of evolution, which are necessary for the computation of divergence time from the phylogeny, were retrieved from the output files of Phylobayes. These rates are calculated by the software as described by the developers elsewhere (Kishino et al., 2001; Lepage et al., 2007) and are expressed as amino acid changes per site per unit of time. The rate at node **D0** was termed **ν**_**max**_ and a baseline rate of evolution during the Proterozoic was obtained as the average of all node rates in Group 4 D1 and D2 and denoted **ν**_**min**_. All sequence datasets and estimated divergence times for each node of each tree used in this analysis are provided in the supplementary data files.

